# Antitachycardia Pacing is More Effective When Delivered to the Left Bundle Branch Area Compared to the Right Ventricle in a Pre-clinical Model of Ischemic Ventricular Tachycardia

**DOI:** 10.1101/2025.08.25.672044

**Authors:** Emmanuel Offei, Yuki Ishidoya, Douglas Smego, Sofia Ruiz Castillo, Muhammad S. Khan, Kyoichiro Yazaki, Ankur Shah, Ava Yektaeian Vaziri, Matthias Lange, Annie M. Hirahara, Hui Li, Gregory J. Stoddard, Ravi Ranjan, Derek J. Dosdall

## Abstract

**Background:** Anti-tachycardia pacing (ATP) delivered from implantable cardioverter defibrillators (ICDs) provides critically timed pacing pulses to terminate ventricular tachycardia (VT). Physiological pacing through left bundle branch area (LBBA) pacing has emerged as a clinically relevant alternative to induce synchronous activation of the ventricles. The main objective of this study was to compare the efficacy and safety of ATP delivered to an LBBA lead and a conventional RV lead.

**Methods:** Using a preclinical animal model (n=7), pacing leads were implanted in the RV apex and LBBA and connected to ICDs. The left anterior descending artery was occluded for two hours to cause an ischemia-reperfusion injury (IRI). Four days following IRI, VT episodes were induced using programmed electrical stimulation (PES), and burst ATP therapy was delivered to the RV or LBBA leads for each VT episode.

**Results:** VT was induced 80 times, with a mean VT cycle length (VT CL) of 180.0±30.0 ms. ATP delivered to the LBBA terminated VT more often than RV ATP (70.2% vs 47.3%, P = 0.04), and there was no significant difference in negative outcomes of VT acceleration or VF induction. Activation sequences from the basket catheter were determined for the pre-therapy VT and captured ATP beats, and correlation coefficients were computed to the activation sequences during ATP delivery. The number of ATP pulses required for the activation sequences to correlate with the captured ATP pattern rather than the pre-therapy VT pattern was lower for LBBA ATP compared to RV ATP (4.1 vs. 5.1 beats, P = 0.04). Successful LBBA ATP demonstrated earlier and more frequent Purkinje activations preceding myocardial activation than successful RV ATP.

**Conclusion:** Improved performance of LBBA ATP compared to RV ATP provides further incentive for the wide use of LBBA leads in patients who need cardiac electrotherapy.

## INTRODUCTION

Acute myocardial infarction (AMI) occurs once every 40 seconds in the United States, and there is an estimated yearly incidence of roughly 605,000 new cases and 200,000 recurrent cases ^1^. The onset of AMI triggers a rapid change in the electrophysiological properties of the ventricular myocardium. In many patients, this causes electrical disturbances in the activation and repolarization of the myocardium that lead to ventricular tachycardia (VT) or ventricular fibrillation (VF). Many cases of sudden cardiac arrest (SCA) and sudden cardiac death (SCD) are caused by ventricular tachycardia and ventricular fibrillation, as the heart is unable to maintain hemodynamic stability. Globally, the incidence of out-of-hospital cardiac arrest (OHCA) is estimated to be 55 per 100,000 persons each year, and the survival rates range from 2% to 11% ^2^. Cardiovascular diseases cause nearly 20.5 million deaths across the world each year ^3^. Out of this number, SCDs account for 4-5 million deaths globally, and in the United States alone, an estimated 300,000 deaths occur annually as a result of SCDs^4^.

Antiarrhythmic drugs can change the electrophysiological properties of the reentrant circuit and can suppress potential triggers for the genesis of VT. Reports have, however, revealed that about 40% of patients receiving antiarrhythmic drugs for sustained VT will receive recurrences within 2 years ^5^, and a VT recurrence can be life-threatening ^6^. Implantable cardioverter defibrillators are superior to antiarrhythmic drugs in improving survival rates after a VT episode and preventing SCD ^7–9^. Thus, patients who are at high risk for SCD may be fitted with an implantable cardioverter defibrillator (ICD). About 800,000 people have ICD implants in the United States alone, and roughly 150,000 new ICD implantations are done annually ^10^. ICDs include an anti-tachycardia pacing (ATP) algorithm that generates critically timed low-energy pulses to terminate VT. If ATP fails in the treatment, high-energy shocks are delivered from the device to abort the VT. These high-energy defibrillation shocks may, however, cause damage to cardiac tissue ^11^, higher mortality ^12^, and lead to increased psychological distress, anxiety, and depression in patients receiving such shocks than in patients who receive no shocks ^13, 14^. Conventionally, ICD leads are implanted at the right ventricular apex, and there is a report that ATP delivery from this location can only terminate about 52%-72% of fast VT ^15^ and has also been associated with adverse therapeutic outcomes such as VT acceleration or degeneration to VF. The problems resulting from defibrillation shocks delivered after ATP fails make an alternate pacing location within the heart, other than the right ventricular apex, essential.

Conduction system pacing delivered to the left bundle branch area (LBBA) has been shown to improve ventricular synchronicity ^16–18^. It shows a survival benefit compared to standard right ventricular or biventricular pacing ^19^. We recently demonstrated that ATP delivered to the His bundle led to fewer adverse outcomes than right ventricular (RV) ATP^20^. With the work presented here, we hypothesize that ATP delivered at the LBBA will lead to increased capture of the ventricular myocardium during VT and thus improved ATP efficacy compared to traditional RV ATP pacing. As a secondary outcome, we hypothesize that LBBA ATP would lead to fewer incidents of VT acceleration and degeneration to VF than RV ATP.

## METHODS

The data that support the findings of this study are available from the corresponding author upon reasonable request.

The experimental protocol conforms to the Guide for the Care and Use of Laboratory Animals and was approved by the Institutional Animal Care and Use Committee (IACUC) of the University of Utah.

### Lead and ICD implantation

Normal mongrel dogs (n=7, 30.0±2.1 kg, 1 year old) were obtained from a Class A vendor and were quarantined for a minimum of 1 week prior to enrolment in the study. Our sample size was determined by our previous experience with this ischemia-reperfusion injury model^20, 21^. Subjects were pre-anesthetized with fentanyl (2-10 mcg/kg, IV) and then anesthetized with Propofol (5-8 mg/kg, IV). Anesthesia was maintained with isoflurane (1.0-3.0%, delivered in 50% oxygen/ 50% room air) through a ventilator in a dedicated operating room for survival large animal surgery. The heart rate, body temperature, blood oxygen saturation, and pCO_2_ were monitored and maintained within normal physiological ranges. A 12-lead ECG was recorded and evaluated during the study using PowerLab (ADInstruments, Dunedin, New Zealand).. A constant rate injection (CRI) of fentanyl was given at 2-10 mcg/(kg.hr), and Cefazolin (20 mg/ kg, IV) was administered every 90-120 minutes during the study. In the event of intraoperative hypotension (a mean arterial pressure < 60 mmHg), a bolus of Lactated Ringer’s solution (75-200 mL, IV) was administered. If hypotension persisted, a dopamine CRI infusion was given 2-10 mcg/(kg.min). If sinus bradycardia developed, atropine (0.01-0.05 mg/kg, IV bolus) was administered to restore regular heart rates.

For lead implantation, a jugular vein was exposed, and under fluoroscopic imaging, two leads were implanted at the RV apex (5076 CapSureFix Novus MRITM, Medtronic, Inc.) and the LBBA (3830 SelectSecureTM, Medtronic, Inc.) (Figure 1). LBBA and RV capture were confirmed by evaluating the transitions in the paced QRS morphology from ECG leads V1 and V6. For LBBA capture, we noticed a narrow QRS complex and a similar wave morphology to sinus rhythm. The QRS complex during RV pacing was broad, and the morphology was characteristically different from sinus rhythm. Once lead placement was confirmed, a subcutaneous pocket was created adjacent to the jugular access site. The leads were then attached to two ICDs (CobaltTM XT DR, Medtronic Inc.), which were implanted into the pocket.

**Figure 1.**
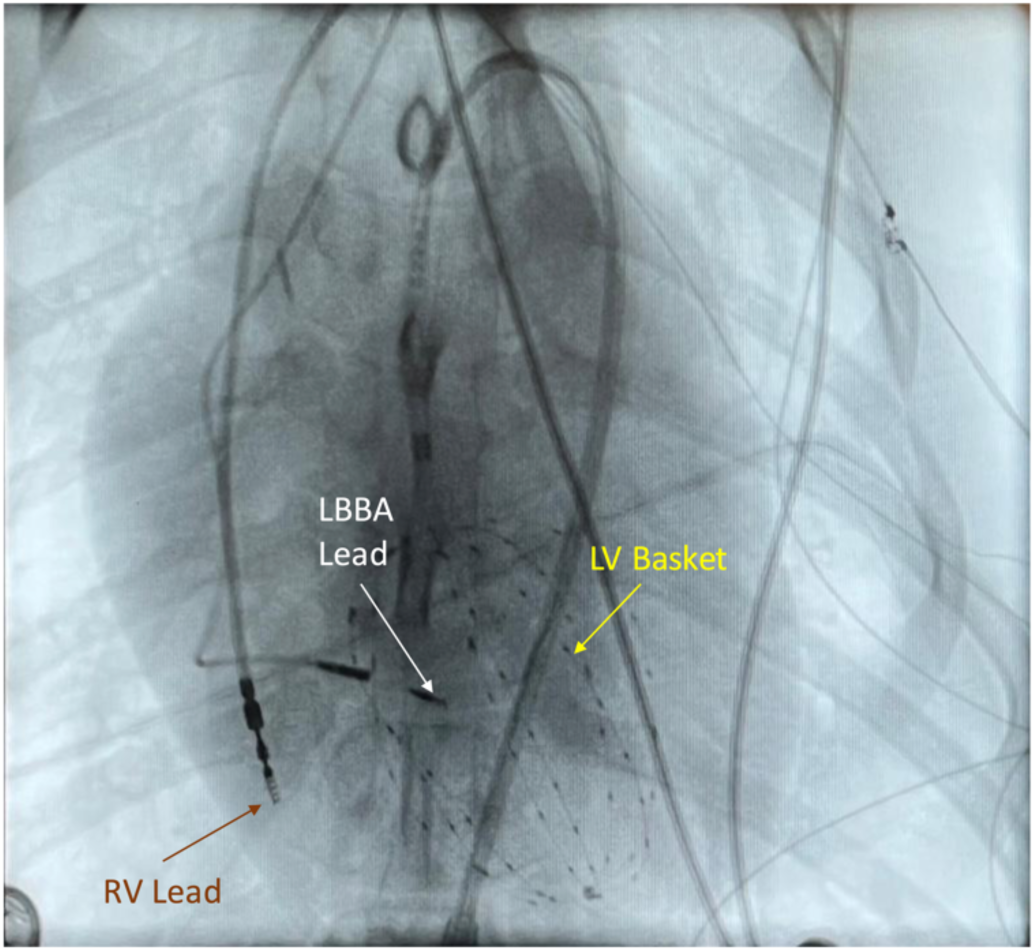
Lead and basket implantation. Example of a fluoroscopic projection (anterior-posterior view) showing the LV endocardial basket (yellow arrow) and the pacing leads implanted at the RV apex (brown arrow) and the LBBA (white arrow).

### Ischemia-reperfusion injury

After completion of the lead and ICD implantation, a left anterior descending artery IRI was induced, as described previously ^22^. In brief, a lateral thoracotomy was performed, and the proximal and distal left anterior descending artery was occluded for 2 hours. After the release of the occlusions, the thoracotomy site was closed, and the animals were recovered from anesthesia.

### Terminal study

As previously described in ^22^, the animals were anesthetized again on the fourth day following the IRI (with the same procedure as above). Twelve (12) - lead ECG recording was implemented, and pacing thresholds for the LBBA and RV leads were confirmed. A 64-channel endocardial basket (FIRMap AR064050, Abbot) was placed in the left ventricle from the femoral artery via a retroaortal approach to record the endocardial electrical activity (ActiveTwo, BioSemi, Amsterdam, Netherlands) during VT induction and ATP therapy. VT was induced using two electrodes from the endocardial basket through PES. If sustained VT was not achieved with programmed pacing, lidocaine (2 mg/kg, bolus) and/or flecainide (50 mg/kg/min, CRI, delivered in a 5% dextrose solution) were administered, and programmed pacing was used to induce VT. When monomorphic VT was detected and sustained for at least 20 seconds, the burst ATP was delivered to the LBBA or RV lead. The order of LBBA or RV pacing was determined randomly, with an effort to balance the number of trials for both pacing locations within each animal. Eight (8) ATP pulses were delivered at 88% of the VT CL in trains until VT was terminated. If VT was not terminated after the fifth ATP train or if VT degenerated to VF, a transthoracic defibrillation shock was delivered. Animals were allowed to recover for at least 5 minutes or until hemodynamically stable. Additional VT episodes were induced and terminated until VT was no longer inducible or the animal did not regain hemodynamic stability.

### Data Analysis

Electrograms from the endocardial recordings were analyzed using custom software developed in MATLAB (MathWorks Inc., MA, USA). A low-pass filter with a 60 Hz notch filter was applied to the recordings to remove noise from the raw electrograms and their temporal derivatives. Five consecutive VT beats were analyzed for each VT episode before ATP therapy was started. The beats when the last train of ATP pulses was applied were also analyzed (ATP beats). The activation time for each beat was determined for all the electrodes using the minimum temporal derivative obtained from the raw electrograms. The activation time was picked manually whenever the algorithm failed to find the most negative dv/dt of the raw signals. Activation maps were created from the activation times. The characteristic capture pattern for the LBBA and RV pacing for each animal was selected from the activation maps (Figures 2A & 2B). For each animal, the beats for the LBBA and the RV captured patterns from successful ATP episodes, had activation sequences that emerged when the last few ATP pulses were applied, and were common for successful ATP trials.

**Figure 2.**
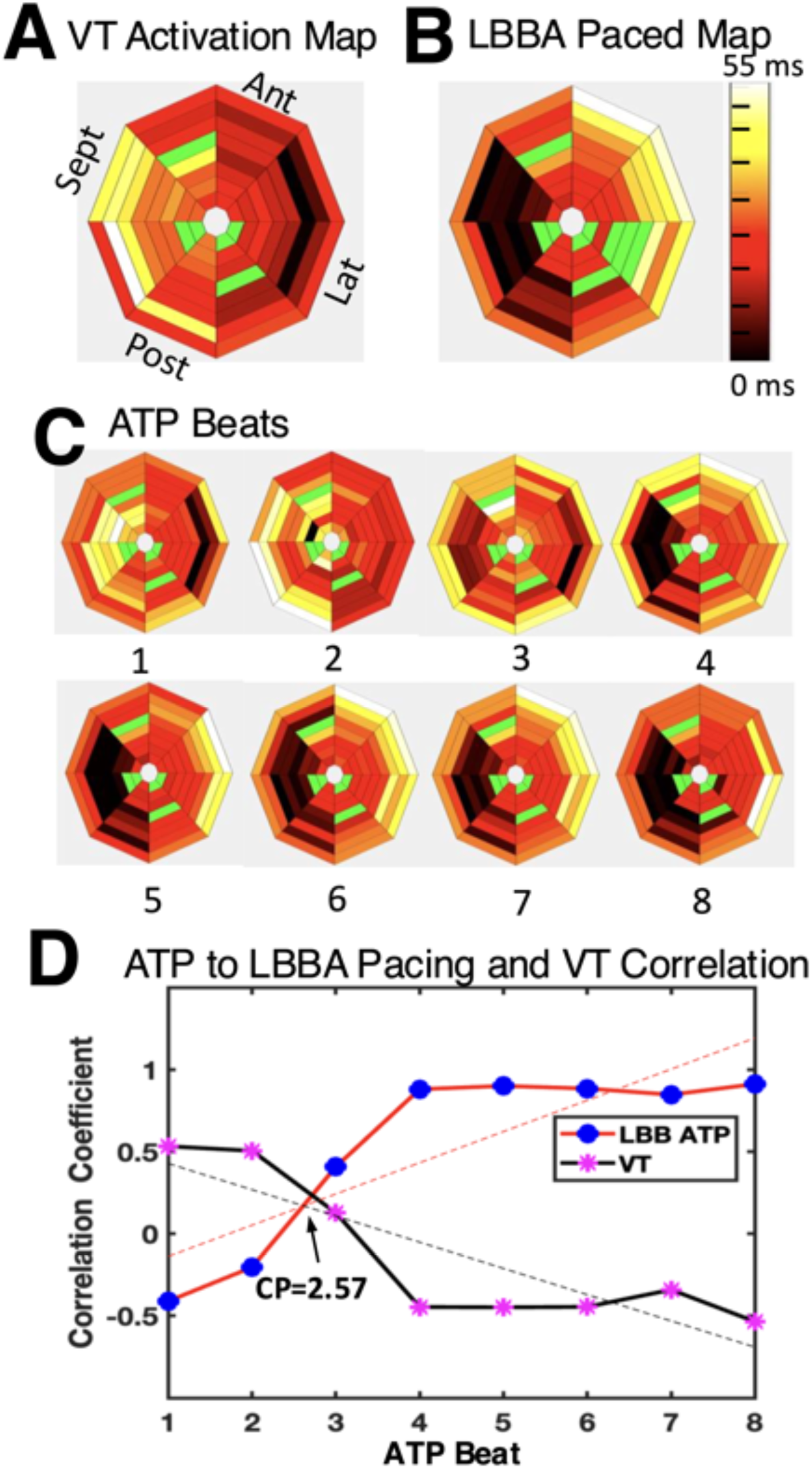
LV activation maps and correlation coefficients. An example of an LV endocardial basket activation sequence for a VT beat (**A**), a captured LBBA pacing activation sequence (**B**), and each of the 8 ATP activation sequences (**C**) for a VT episode treated with LBBA ATP. Bad electrodes are marked in green. The center of each activation plot shows LV apical basket electrodes, and the outer ring shows electrodes near the LV base. Spearman’s correlation coefficients are plotted (**D**) for of each of the 8 ATP activation sequences to the VT activation sequence (**A** vs. each activation sequence from **C**, solid black line with red asterisks in **D**) and to the LBBA captured activation sequence (**B** vs. each activation sequence from **C**, solid red line with blue circles in **D**). The dashed lines show the linear fits for 8 Spearman’s correlation coefficients for both the ATP (red) and VT (black) activation sequences. The point at which the linear fit of the ATP activation sequence correlates more with the captured LBBA activation sequence than the VT pattern is the crossover point (CP).

Purkinje activations were determined through bipoles created by subtracting each unipolar signal from the neighboring unipolar signal on each spline in MATLAB. From the 64 unipolar basket electrodes, 56 bipoles were formed, and channels with noisy electrograms were omitted. The His-Purkinje activations were found during sinus rhythm, the progression of the VT wavefronts, and finally, at the time of ATP delivery. As previously reported, Purkinje activations were identified based on their morphology, timing, and amplitude^23^. Purkinje activations were identified manually and marked as rapid, short-duration activations (about 1-2 ms). Time intervals from the end of the last ATP pulse to the earliest Purkinje and myocardial activations were measured for successful ATP trials. The number of these was 44.

### Statistical Analysis

Success rates for terminating VT with ATP delivered from the RV and LBBA leads were compared with a mixed-effects logistic regression model in Stata 17.0 (StataCorp, College Station, TX, USA). This model accounted for repeated measures and varying numbers of trials in each animal. Percentages and standard errors for each pacing location were obtained from a post-fit marginal estimation.

The correlation coefficients between the ATP beats (Figure 2C) and the activation sequences from the LBBA and RV capture patterns were calculated using Spearman’s Rank Correlation test, MATLAB (MathWorks, B, Natick, MA, USA) (ATP coefficients). The same analysis was done between the ATP beats and the VT beat for each VT episode (VT coefficients). A linear regression was fitted to the ATP and the VT coefficients for each VT episode, and the intersection point (termed crossover point) for the two lines of best fit was identified (Figure 2D). A crossover point of 1 was assigned for linear regression intersections < 1, and a crossover point of 8 was assigned for the linear fits’ intersections of>8. A mixed effects linear regression model was created in Stata to compare the slope of the ATP and the VT coefficients linear regressions for the LBBA and RV ATP episodes.

## RESULTS

### Therapeutic Efficacy of ATP

One out of the seven dogs initially enrolled in the study was eliminated because we failed to resuscitate the dog from a VF event during the terminal study. This exclusion criterion was determined *a priori*. From the surface ECG recordings, the QRS duration during LBBA capture was narrower than that for RV capture (70.6±8.2 ms vs. 111.0±11.4 ms, P = 4×10^-4^). The QRS durations were measured during sinus rhythm, and at the time when we paced both the LBBA and the RV to determine their capture threshold. Eighty (80) episodes of VT were induced. Still, one episode of VT was eliminated from the analysis because competing ATP pulses were delivered from the RV and LBBA pacing locations during the ATP trial. The mean VT CL for all the analyzed episodes was, 180.0±30.0 ms. The VT CL was not significantly different between the episodes for the LBBA and the RV. Out of the 79 VT episodes that were analyzed, 41 ATP trials were made at the LBB, and the RV was tested with 38 ATP trials. Out of these trials, LBBA had 91 trains of ATP pulses, and the RV had 116 trains of ATP pulses. An overall success rate of 65% was recorded for ATP delivered at both pacing sites. Of all the successful cases, 73% were achieved with the first train of ATP pulses and 24% were achieved with the second train of pulses..

The resumption of sinus rhythm characterized successful events after the last ATP pulse was applied (Figure 3A), and unsuccessful events had ongoing VT wavefronts after the last ATP pulse (Figure 3B). ATP trials at the LBBA were successful 70.2±10.2% of the time, while 47.3±11.3% of the trials at the RV were successful (RR = 1.44, 95% CI (1.03,2.01), P = 0.04) (Figure 3C). Observations of a few or several extra VT beats following the last ATP pulse were made for the successful events. Successes with no extra VT beats were classified as Type A successes, and those with one or more extra beats as Type B successes. ATP delivery to the LBBA resulted in a higher percentage of Type A successes than ATP delivery to the RV (26.3% vs. 18.4%). There was also a higher percentage of Type B successes for ATP delivery to the LBBA than the RV (55.3% vs. 34.2%). However, there was no statistically significant difference in the incidence of Type A and Type B successes between the pacing sites (P = 0.99).

**Figure 3.**
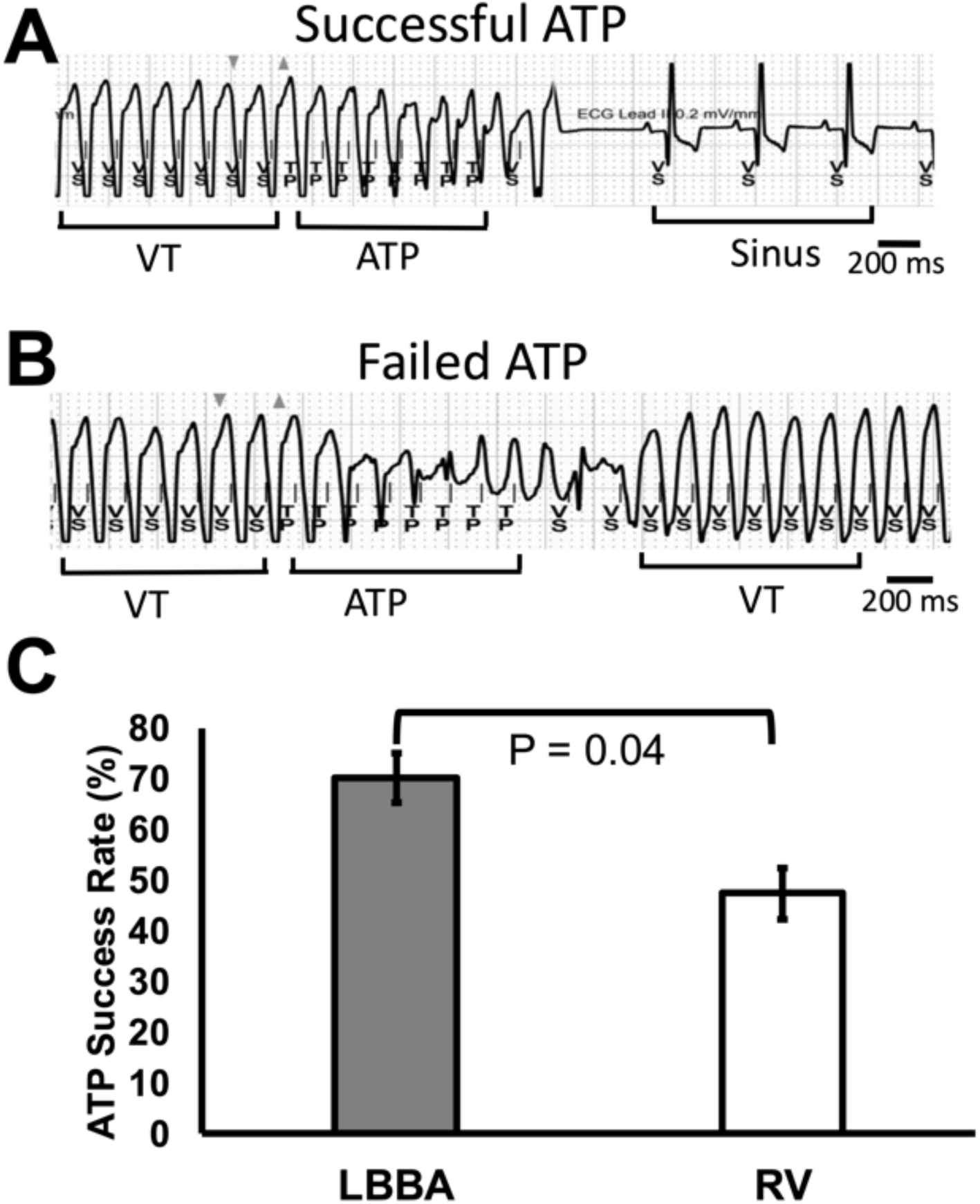
ATP therapy outcomes. During an example VT episode, ATP successfully terminated VT from an LBBA lead (**A**). ATP delivered from an RV lead failed to terminate VT in another episode (**B**). Out of all ATP trials (n=79), LBBA ATP showed an increased success rate compared to RV ATP (**C**). (n=6 mongrels). TP = Tachycardia Pacing, and VS = Ventricular Sense. Success rates were compared using a mixed-effects regression model.

### ATP Negative Outcomes

Negative outcomes associated with ATP delivery were observed for both pacing sites. There were incidents of VT acceleration and the degeneration of VT to VF. The VT CL decreased by 10% or more for VT acceleration after the last train of ATP pulses (Figure 4A), and the ECG waveforms looked chaotic and disorganized with no identifiable P waves, QRS complexes, or T waves for VT events that degenerated to VF after the last train of ATP pulses (Figure 4B). The mean CL for all negative events was 170.0±30.0 ms. An overall negative outcome of 12.7% was recorded for ATP delivered to both pacing sites. LBBA ATP was accompanied by more negative outcomes than RV ATP (17.0±*7.9* % vs. 11.7 ± 6.5%, P = 0.51) (Figure 4C), although this difference failed to reach statistical significance.

**Figure 4.**
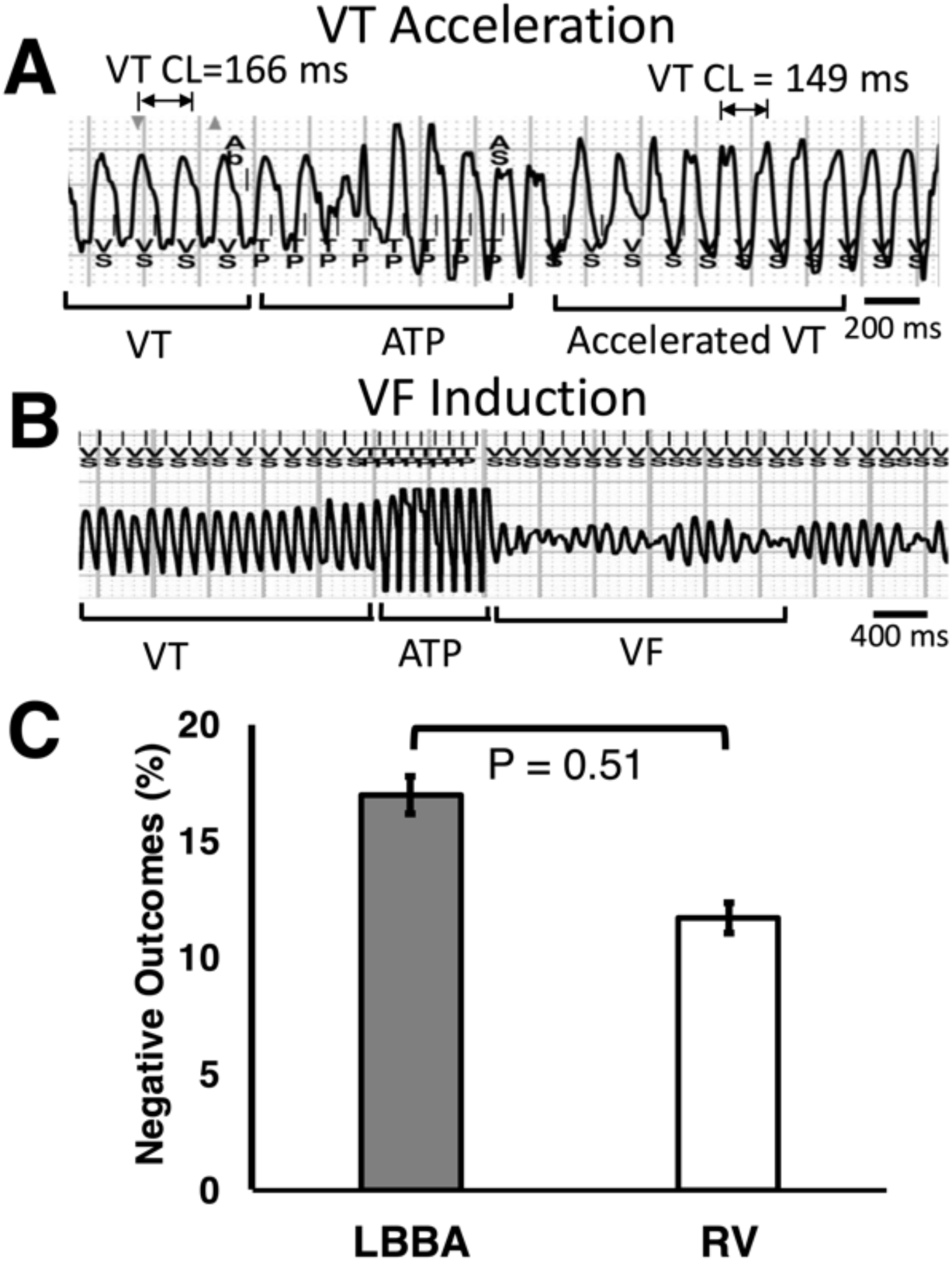
ATP negative outcomes. During an example VT episode, ATP delivered to an RV lead caused the VT cycle length to be accelerated (**A**). VT degenerated to VF after ATP was delivered to an RV lead in another VT episode (**B**). For all ATP trials (n=79), negative outcomes for LBBA vs. RV ATP were not significantly different (C). (n=6 mongrels).VT CL = VT Cycle Length, TP = Tachycardia Pacing, and VS = Ventricular Sense. Safety of ATP was compared with a mixed-effects regression model.

### Myocardial Capture with ATP

For this analysis, only 70 out of the 79 endocardial recordings were used. Eight VT episodes were excluded from the analysis because the signal-to-noise ratios of the electrograms from all the channels were insufficient to accurately identify activation times and sequences confidently. One additional VT episode was also excluded because we had no recording of the pre-ATP VT and the ATP beat for the first ATP pulse.

From the 70 endocardial recordings that were used for this analysis, LBBA ATP had 37 ATP trials, and RV ATP had 33 ATP trials. For both pacing locations, ATP coefficients showed low correlation between the first few ATP beats and the LBBA/ RV capture pattern, but that increased as more ATP beats were added (Figures 5A & 5B). In contradistinction to this, for both pacing sites, VT coefficients for the first few ATP beats showed high correlation with the VT beats, but that correlation declined as more ATP pulses were delivered (Figures 5C & 5D). A statistically significant positive effect existed between the pacing site and the slope of the lines of best fit, for the ATP coefficients. Changing from the RV to the LBBA for ATP delivery increased the slope by approximately 0.045 units (P = 6.0×10^−4^). Switching from the RV to the LBBA for ATP delivery decreased the slope of the best fit lines for the VT to ATP pattern by approximately −0.055 units (P = 3.6×10^−5^). The crossover point marked the point in time after which the ATP coefficients became larger in magnitude than the VT coefficients, and it represented a disruption to the VT circuit. Generally, ATP to the LBBA had a crossover point, one full beat earlier than the crossover point for ATP delivery to the RV (4.1 vs. 5.1, P = 0.04) (Figure 6A). The crossover point for the overall ATP outcome was shorter for successful vs. failed ATP (3.7 vs. 5.6 beats, P = 1.0×10^−4^, Figure 6B). Additionally, for both pacing locations, the crossover point for all successful cases was shorter in duration than that for failed outcomes. For the LBBA, the number of beats was 3.3 vs. 5.81 beats (P = 0.001, successful vs. failure, Figure 6C), whereas the crossover point for the RV was 4.0 vs. 5.7 (P = 0.003, successful vs. failure, Figure 6D). For successful ATP trials, LBBA ATP had a crossover point that was 0.87 beats earlier than RV ATP, but this trend did not achieve statistical significance (P = 0.08, Figure 6E). The interaction between the pacing site and unsuccessful events had no statistical significance (P = 0.70, Figure 6F).

**Figure 5.**
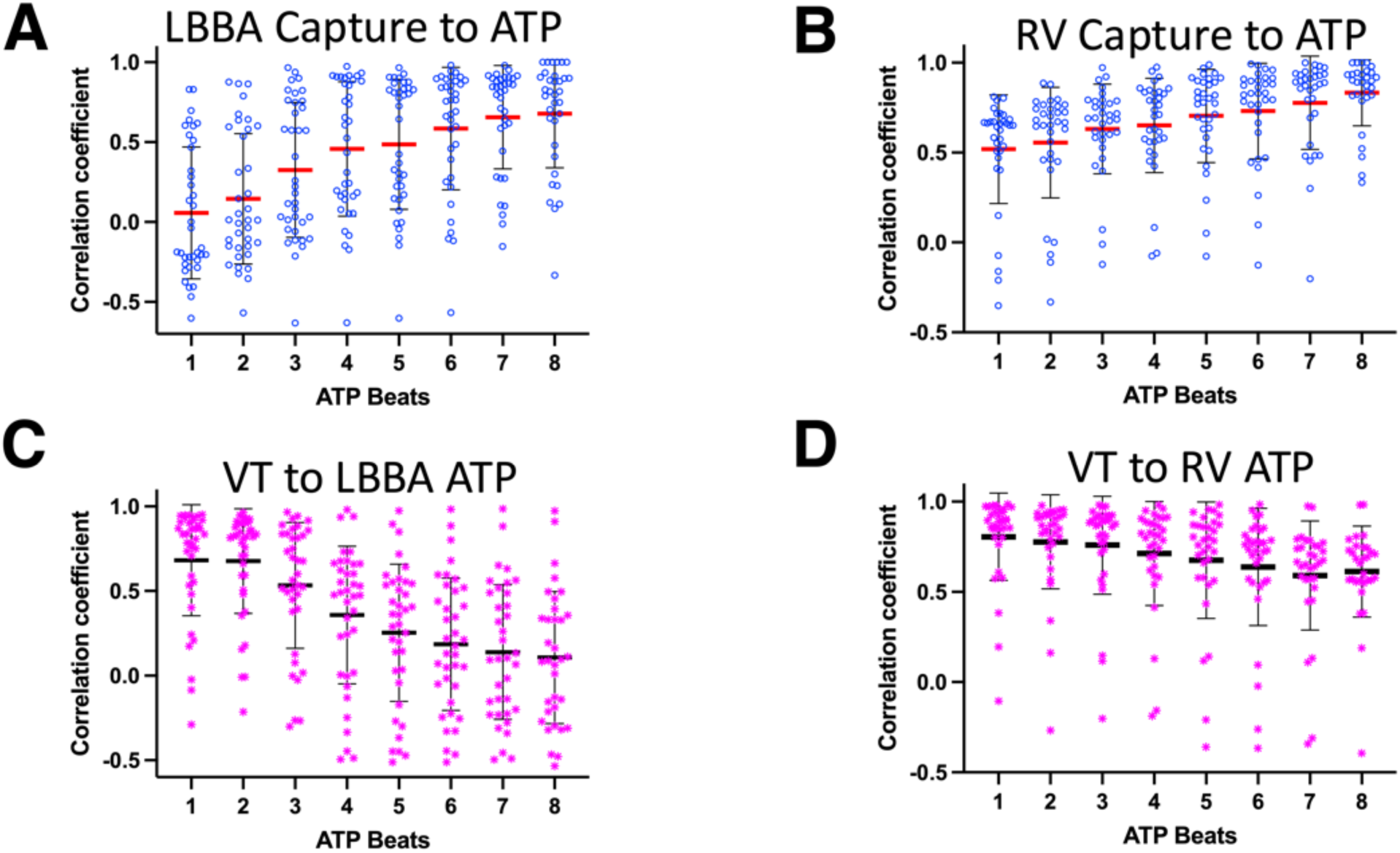
Activation sequences during ATP delivery. Spearman’s Rank correlation coefficients for the activation sequences during ATP. For LBBA ATP trials (n=37) (**A**) and RV ATP trials (n=33) (**B**), the mean ATP-captured correlation coefficients (red lines) increased as more ATP pulses were delivered. The mean correlation coefficients (black lines) of the VT activation sequence decreased as more ATP pulses were applied to the LBBA (**C**) and the RV (**D**). Coefficients for the ATP-captured activation sequences are marked with blue circles, and the coefficients for the VT activation sequences are marked with magenta asterisks. (n=6 mongrels).

**Figure 6.**
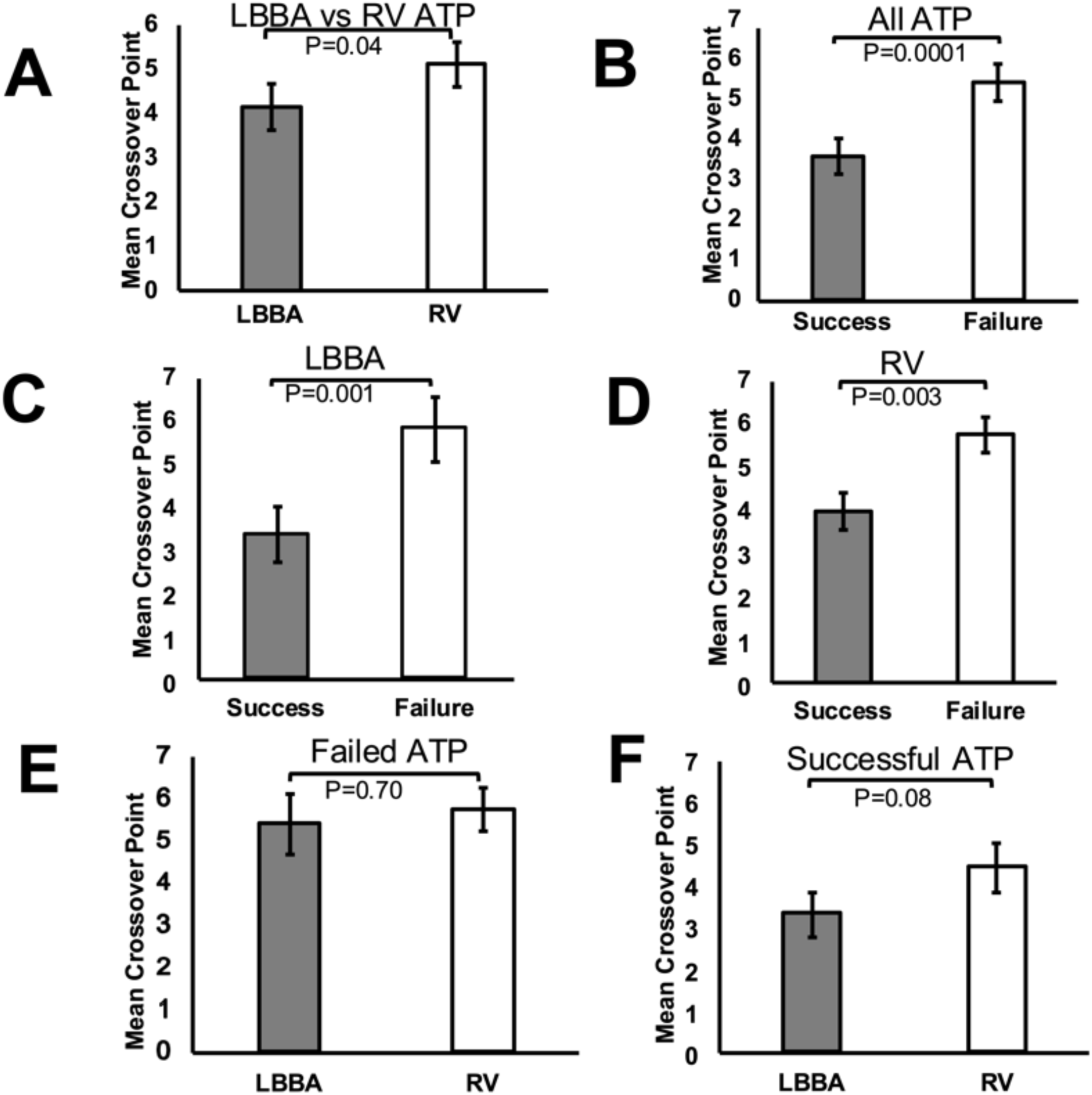
Crossover points as a marker for VT circuit disruption. For all ATP trials (n=70) used in the analysis, the crossover points between the correlation coefficients of ATP captured beats and VT beats during ATP therapy were one full beat earlier for the LBBA than the RV (4.1 vs. 5.1, respectively) (**A**). The crossover points were nearly two beats earlier for successful vs. failed trials (3.7 vs. 5.6, respectively) (**B**). For LBBA ATP trials (n=37), the successful trials had a crossover point that was 2.5 beats earlier than the failed trials (3.3 vs. 5.8, respectively) (**C**). Whereas for the RV ATP trials (n=33), the successful trials were 1.8 beats earlier than the failed trials (4.0 vs. 5.8, respectively) (**D**). For failed ATP trials (n=26), LBBA had a lower crossover point compared to RV ATP (5.4 vs 5.7, respectively) (**E**). In successful ATP trials (n=44), the LBBA crossover point was earlier than the RV crossover point (3.3 vs 4.2, respectively) (**F**). A mixed-effects regression model was used to evaluate the effect of the crossover point on the different outcomes. (n=6 mongrels).

### Timing for Earliest Purkinje Fiber and Working Myocardium (WM) Activation

On average, Purkinje activations were detected on 20 of the 56 bipoles created from the 64 unipoles of the basket catheter. Activations spread through the Purkinje fibers into the WM during both sinus rhythm and VT, as the Purkinje activations were seen to precede each WM activation during VT (Figures 7A & 7B). During ATP delivery that successfully terminated VT, Purkinje activations were detected earlier following the final ATP pulse delivered to the LBBA than when it was delivered at the RV (18.8±5.4 ms vs. 39.8±5.6 ms, P = 2.1×10^−11^, Figure 7C). Similarly, myocardial activations were seen to be earlier when ATP was delivered to the LBBA than when it was delivered to the RV (25.7 ± 5.1 ms vs. 38.7 ± 5.4 ms, P = 2.0×10^−4^, Figure 7D). Purkinje activations were detected earlier than myocardial activations more frequently for the LBBA ATP than for the RV ATP therapy (84.7 ± 10.7 % vs. 41.8 ± 8.3 %, P = 8.0×10^−4^, Figure 7E).

**Figure 7.**
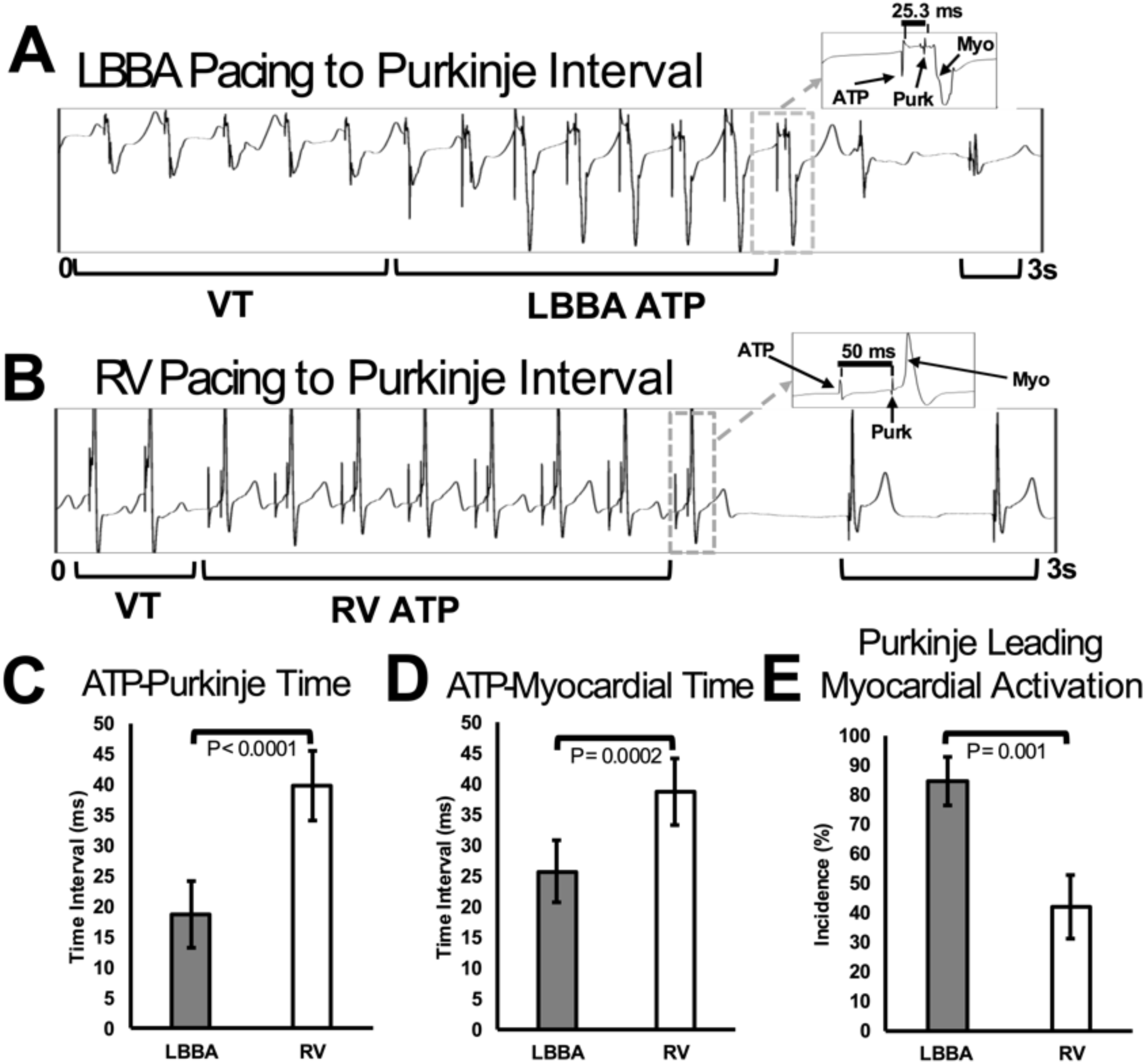
Capture of Purkinje fibers and working myocardium during successful ATP delivery. Electrograms from a single channel of the endocardial basket for an LBBA ATP episode (**A**) and an RV ATP episode (**B**) show VT wavefronts, ATP delivery, and restoration of sinus rhythm. Local Purkinje activations can be seen before each myocardial activation, as shown in the expanded last ATP beats in (**A**) and (**B**). Out of all successful ATP trials (n=44), ATP delivery to the LBBA caused the Purkinje fibers to be activated earlier than when ATP was delivered to the RV (**C**). The WM was also activated earlier when ATP was delivered to the LBBA than when it was delivered to the RV (**D**). Activation of the Purkinje fibers occurred before the WM more often when ATP was delivered to the LBBA compared to the RV (**E**). Myo = myocardial activation, and Purk = Purkinje activation. A mixed-effects regression model was used to evaluate the timing of the Purkinje and WM activations when ATP was delivered to the LBBA and the RV. (n=6 mongrels).

## DISCUSSION

This study compared the safety and therapeutic efficacy of ATP delivered in the RV to the LBBA. The findings in this work include: 1) LBBA ATP successfully terminated VT with a higher success rate compared to RV ATP, 2) negative outcome rates of VT acceleration and VF induction from unsuccessful ATP delivery were not significantly different between LBBA and RV ATP, 3) LBBA ATP disrupts the VT activation pattern and captures the activation sequence in fewer beats than RV ATP, and 4) successful ATP delivered to the LBBA had a shorter pacing to Purkinje interval than RV ATP, suggesting that LBBA ATP spreads through the conduction system. These findings provide valuable evidence supporting the use of LBBA ATP in a relevant pre-clinical VT model.

Physiological pacing through a lead in the LBBA has emerged as a promising therapy that leads to shorter QRS durations, less intraventricular dyssynchrony, and improved clinical outcomes compared to traditional RV pacing ^24–26^. The narrower QRS duration observed in this study during LBBA capture compared to RV capture highlights the more physiologic activation of the ventricles. The increased incidence and shorter time gap observed following successful LBBA ATP therapy is consistent with LBBA ATP activating the Purkinje network for more extensive and synchronous myocardial activation, leading to improved ATP success rates for LBBA compared to RV ATP.

Only a paucity of information is available in the literature on treating ventricular arrhythmias with ATP or shock therapy delivery through an LBBA lead. Clementy et al. showed the feasibility of a lead implanted in the LBBA to detect induced VF successfully and terminated that with a separate RV ICD lead^27^. Recent studies have demonstrated that LBBA pacing leads with defibrillation shocking coils can be successfully implanted and have been used to terminate induced VF ^28, 29^. These studies demonstrate a clinical interest in using LBBA leads not only for pacing therapy but also for detecting and terminating dangerous ventricular arrhythmias.

Moreover, our findings on the negative outcomes of ATP therapy may be explained by the nature of the VT CL, the mean variation in VT CL, and the similar VT morphologies for both the LBBA and RV ^30^. There was a non-significant increase in negative outcomes following unsuccessful LBBA vs. RV ATP. This difference may be attributed to the temporal proximity of the stimulation of the LBBA, and the activation of the Purkinje system, where precise timing is critical to avoid unintended entrainment of the reentrant circuit ^31^. VT acceleration can occur when ATP fails to reach the excitable gap at the right moment in time of the ongoing VT wavefront and can, therefore, entrain it. This may occur particularly in fast VTs, where the reentrant circuit’s excitable gap is reduced in duration, and the ATP pulses cannot interact with the circuit at the right time for VT termination. Rather than terminating the reentrant circuit, LBBA ATP may entrain the myocardium, resulting in acceleration of the circuit. The mean CL for the negative events (170.0 ± 30.0 ms) was 5.8% less than the mean CL for all the VT episodes 180.0±30.0 ms. Therefore, the excitable gap was reduced for these events, and the ATP pulses may have had a greater probability to entrain the VT circuit and myocardium. From the 12-lead ECG electrogram data, we recorded no statistical significance between the VT CL for LBBA and RV ATP, and the VT morphologies were similar. Our finding differs from the results we gathered from our earlier work on the safety and efficacy of RV ATP vs. His-bundle ATP ^20^. His bundle ATP led to significantly fewer negative outcomes than RV ATP. The difference in ATP safety between His and LBBA compared to RV ATP may be attributable to the overall more interventricular synchrony of His bundle pacing compared to LBBA pacing or methodological differences (His study conducted with external pacer compared to ICDs in LBBA study).

The comparable VT CLs observed between the VT episodes for LBBA and RV pacing indicate that the intrinsic properties of the VT were not significantly different for therapy delivered to the two pacing sites. The differences in the therapeutic outcomes for LBBA ATP and RV ATP can, therefore, be attributed to the pacing location rather than the variations in the reentrant VT circuit itself^32^.

A significant observation was that most successful ATP events, irrespective of the pacing site, occurred after the first train of ATP pulses was delivered (73%). This finding underscores the importance of early intervention and the efficacy of the initial ATP trials in terminating VT. Wathen et al. reported that 85% of fast VT episodes were terminated with the first train of ATP pulses ^33^. The apparent disparity in the two results can be attributed to the differences in the cycle lengths of the VT episodes from both studies. VT CL in our study (180±30 ms mean) were faster than their reported results (301±24 ms). Termination of VT is more difficult when the VT CL is reduced ^34–36^. Also, ATP delivered to the conduction system during VF has been shown to intermittently cause activation patterns similar to sinus rhythm, but ATP methods have not been successful in consistently terminating VF ^37^.

The results on the dynamics of ATP and VT coefficients during ATP reveal differences in the influence pacing sites have on these coefficients, which sheds some light on the mechanisms that underlie the therapeutic efficacy of ATP. Entrainment of the WM during VT through the pacing of the LBBA engaged the conduction system effectively and caused the WM to be captured physiologically. There was a progressive entrainment of the reentrant VT circuit that was greater for LBBA ATP than RV ATP. The slope of the line of best fit to the ATP coefficients for LBBA ATP was steeper than the slope for RV ATP. This implies that the buildup of pacing effectiveness for LBBA ATP is accelerated compared to RV ATP. Additionally, the reduction in the slope of the line of best fit to the VT coefficients when the pacing site is changed from the RV to the LBBA also suggests a more rapid attenuation of the reentrant VT circuit during LBBA ATP. In particular, during LBBA ATP, the train of ATP pulses spreads rapidly and has greater access to the reentrant circuit for collision to occur, or it can overdrive the VT circuit to a higher degree to reset it. This postulation stems from the fact that pacing the LBBA is more physiologic, as the entrainment of the WM is mediated through the intrinsic conduction system. There was, therefore, a more synchronous activation of the ventricles, as was validated by the narrow QRS complex during the LBBA pacing we observed. Entrainment of the WM during RV ATP is not as rapid because the pacing occurs outside the natural conduction system ^38^. The abnormal electrical activation progresses through the ventricular muscle, and it yields a delayed activation of the WM that is made evident by the prolonged QRS duration ^39, 40^ during RV ATP. Our findings agree with those in the existing literature, where better electrical synchrony and, thus, more rapid WM entrainment are reported for LBBA pacing than RV pacing ^41–44^.

A key finding from the work was identifying the crossover point – the point after which ATP coefficients exceeded VT coefficients, representing a disruption of the VT circuit. The crossover point was one full beat earlier when ATP was delivered to the LBBA than when it was delivered to the RV. The results from the crossover point out the fact that depolarization wavefronts spread rapidly from the LBBA to capture the WM. The earlier crossover point for LBBA ATP is consistent with the superior synchronized activation of the WM for ATP delivery to the LBBA. VT termination is typically achieved through resetting, where the refractory region of the VT circuit is shifted close to the stimulation site so that the orthodromic wavefront is blocked ^45^. Since delivering ATP pulses through the LBBA to reach the site of the reentrant VT circuit appears to be favored spatially and temporally over the RV, it is more reasonable that LBBA ATP achieved a rapid disruption of the VT circuit than RV ATP. In their experimental work in an anatomically detailed rabbit, Byrd et al. reported that BiV ATP is more effective than conventional RV ATP ^45^. Pacing simultaneously from spatially distributed leads advanced the orthodromic wavefront more into the refractory region of the VT circuit, blocking it and causing the VT to be terminated.

Additionally, Haghjoo et al. confirmed the same findings in a group of heart failure patients who received BiV ICD implantations ^32^. They reported that BiV ATP and LV ATP had greater efficacy than RV ATP. The MIRACLE ICD trial also presented similar findings for BiV ATP vs. RV ATP; BiV ATP had better therapeutic outcomes. In the InSync ICD study, it was also reported that BiV ATP was more successful at treating VT than RV ATP, leading to fewer accelerations for slow and fast VTs^46^. The ADVANCED CRT-D trial reported that BiV ATP is better than RV ATP in patients with ischemic heart disease, but the efficacy was similar in patients with non-ischemic heart disease ^47^. These studies investigated the therapeutic efficacy of pacing from multiple locations in the heart, as we did, and reiteratively, a site that is proximal to the VT circuit not only spatially but temporally, as we have established through the crossover point, seems to offer greater efficacy for VT termination.

Furthermore, the results on the crossover point for the therapeutic outcomes (successes and failures) of each pacing location suggest that an early resetting of the VT circuit may predict ATP success. The temporal dynamics of beat progression during ATP delivery may be an essential marker for determining the likelihood of pacing success. Fewer beats were required for resetting the VT circuit when ATP was successful than when ATP failed. Ventricular capture appears rapid for successful outcomes for both the LBBA and the RV. In contrast, there seems to be some delayed conduction to the VT circuit in the failed ATP trials, which was made evident by the long crossover point we found for those trials. Several studies have shown that pacing the LBBA provides superior electrical resynchronization compared to traditional RV and BiV pacing ^38, 42, 48–51^. These studies made the improved synchronization evident by the narrow-paced QRS duration they reported. This supports the idea of enhanced ventricular activation times for the successful ATP outcomes reported in our work. Although those studies focused on QRS duration, there is some consistency with the underlying principle – more efficient WM activation is reflected in short crossover duration, which is associated with better ATP outcomes. Briefly, the crossover point provides a quantifiable metric for early prediction of ATP success. Future studies developing real-time analysis and feedback are warranted to determine if adaptive ATP using measures derived from crossover point dynamics can improve ATP success rates and clinical outcomes.

The overall success of ATP therapy for both pacing locations was linked to differences in the crossover point. The crossover point for successful VT termination events was 1.9 beats earlier than failed ATP trials (P = 1.0X10^−4^). This finding underscores the importance of rapid circuit disruption for ATP therapy to be effective, as mentioned earlier. Notably, the crossover point was 0.87 beats earlier when ATP was delivered to the LBBA compared to the RV, although the interaction did not yield statistical significance (P = 0.08). The more synchronous electrical activation of the WM during LBBA ATP than the RV ATP, and the electrophysiological properties of the WM may explain this outcome ^52, 53^. For unsuccessful events, no significant difference was observed between the pacing sites (P = 0.70), with the crossover point for LBBA occurring only 0.34 units earlier than RV ATP. These findings suggest that while LBBA pacing advances the timing of VT circuit disruption, its superiority in VT termination appears nuanced and may depend on patient-specific electrophysiological properties.

Also, our results on the temporal dynamics for Purkinje fiber and WM activations provide novel insights into the mechanisms that underlie ATP effectiveness and pinpoint the importance of the pacing site in achieving rapid termination of VT. The results showed that Purkinje activations preceded WM activations during VT and sinus rhythm. This observation confirms the Purkinje fibers’ critical role as the primary conduction pathway for initiating myocardial activations and propagating activation wavefronts during arrhythmic and normal conditions. Of importance, ATP delivery to the LBBA resulted in significantly earlier Purkinje activations (18.8±5.4 ms) than the RV (39.8±5.6 ms). This suggests that LBBA pacing provides more efficient engagement of the Purkinje system, facilitating earlier reentrant circuit disruption. To the best of our knowledge, a direct comparison of the timings for Purkinje and myocardial activations for LBBA vs. RV pacing, as demonstrated in this study, does not exist in the literature. In their work on impulse propagation through the Purkinje system and WM of intact dog hearts, Ben-Haim et al. reported that Purkinje activation times (defined as the time interval from stimulus to Purkinje spike) increased linearly from the pacing site ^54^. The group reported a 6.4 ± 0.5 ms latency for left ventricular apical pacing. Their findings agree with ours because, for the LBBA, the pacing lead is near the left ventricular conduction system, so we would expect early Purkinje activations. For the RV apical site, however, there is an intervening myocardium from the lead before we reach a Purkinje fiber.

Similarly, WM activations were also observed to occur earlier with LBBA pacing compared to RV pacing (25.7±5.1 ms vs. 38.7±5.4 ms, P = 2.0×10^−4^). This earlier activation reflects a more synchronized and effective depolarization of the myocardium, which likely contributes to the superior performance of LBBA ATP in interrupting the VT circuit. These findings are supported by earlier reports that pacing the conduction system can optimize conduction timing and increase the likelihood of ATP success. Other groups have also reported early capture of the conduction system and the WM during the pacing of the LBBA. Zhang et al. confirmed the capture of the left conduction system by reporting a shortening of the stimulus to peak left ventricular activation time during LBBA pacing ^55^. Likewise, Zhu et al. reported a stimulus to left ventricular activation time (LVAT) of 48.70**±**13.67 ms ^56^. In their work in a canine model, Chen et al. also reported a pacing stimulus to left ventricular activation time of 39.67±1.53 ms and an LBB potential to ventricular activation interval of 12.67±1.15 ms during LBBA pacing ^57^. These results imply a rapid capture of the conduction system and the WM during LBBA pacing, which agrees with our findings. The marginal differences between our results and theirs can be attributed to anatomical differences and differences in the experimental setups.

Moreover, we also observed a higher frequency of Purkinje activations preceding myocardial activations when ATP was delivered to the LBBA than when it was delivered to the RV (84.7±10.7% vs. 41.83±8.3%, P = 8.0×10-4). This result underscores the preferential engagement of the Purkinje fibers during LBBA pacing and suggests that LBBA ATP exploits the native conduction system more effectively than RV ATP. By activating the Purkinje fibers first, LBBA ATP ensures a rapid and organized spread of electrical activity to the WM, crucial for disrupting reentrant VT circuits. RV pacing often initiates a slower, less synchronous myocardial activation sequence, which may delay VT termination or reduce the success of ATP therapy 32, 58. LBBA pacing benefits from the anatomical proximity to the Purkinje system, which allows for a more physiologically relevant and rapid activation sequence.

### Limitations

Firstly, all the VT episodes in the work were induced from PES, which is characteristically different from the spontaneous VTs in humans. Spontaneous VTs are easier to terminate ^59^. However, the comparative nature of the work between the LBBA and RV overcomes this drawback. Also, the VT CL in dogs is faster than that in typical human VT episodes. This is another limitation because the arrhythmias in dogs (with a narrow excitable gap) are more difficult to break than in humans. Therefore, the study’s outcome will likely differ in humans, with a significant likelihood of more successful episodes than reported in the current work. Moreover, the anatomical differences across the animals, the depth of lead penetration into the LBB’s septum, the scar’s geometry, and the distance of the pacing lead(s) from the ischemia site(s) might have affected the study’s outcome. The timeline of our VT induction and ATP testing (4 days post-infarction) differs from that of most patients implanted with ICDs.

## CONCLUSION

This study demonstrated that delivering ATP to the LBBA is more efficacious at aborting VT events compared to when ATP is delivered to the conventional RV apical site. The steeper ATP correlation coefficients, earlier crossover points, and earlier and more frequent Purkinje activations reported for LBBA ATP than RV ATP provide a mechanistic understanding of this outcome. The data we have presented provides incentives for scientific investigation into the electrophysiological properties of the conduction system, making it an ideal site for treating arrhythmias. Furthermore, the work is invaluable as conduction system pacing becomes more common in clinical practice. The improved performance of ATP delivered to the LBBA compared to the RV provides further incentive for employing LBBA leads in cardiac resynchronization therapy and ATP delivery.

## Acknowledgements

The authors would like to thank Orvelin Roman for supporting the work with animal care and surgeries.

## Non-standard Abbreviations and Acronyms

AMI: Acute Myocardial Infarction
ATP: Anti-tachycardia Pacing
CRI: Constant Rate Injection
IACUC: Institutional Animal Care and Use Committee
ICD: Implantable Cardioverter Defibrillator
IRI: Ischemia-Reperfusion Injury
LBBA: Left Bundle Branch Area
OHCA: Out-of-Hospital Cardiac Arrest
PES: Programmed Electrical Stimulation
RV: Right Ventricle
SCA: Sudden Cardiac Arrest
SCD: Sudden Cardiac Death
TP: Tachycardia Pacing
VF: Ventricular Fibrillation
VS: Ventricular Sense
VT: Ventricular Tachycardia
VT CL: Ventricular Tachycardia Cycle Length
WM: Working Myocardium

